# Knockouts of positive and negative elements of the circadian clock disrupt photoperiodic diapause induction in the silkworm, *Bombyx mori*

**DOI:** 10.1101/2022.05.16.492110

**Authors:** Hisashi Tobita, Takashi Kiuchi

## Abstract

Diapause is one of the most important traits that have sustained insects to thrive. Insects can prospectively arrest their development and increase environmental stress-resistance by entering diapause. The photoperiod is the signal that indicates insects the proper timing to enter diapause. Circadian clock genes are shown to be involved in photoperiodic diapause induction in various insect species. The silkworm, *Bombyx mori*, enters diapause at the embryonic stage. In bivoltine strains, diapause determination is affected by embryonic temperature and photoperiodic conditions during embryonic and larval stages. Two independent studies showed that knocking out the core clock gene, *period*, perturb photoperiodic diapause induction. However, whether the circadian clock as whole or individual clock genes are responsible for the photoperiodic diapause induction remains unknown. In this study, using CRISPR/Cas9 we knocked out negative (*period* and *timeless*) and positive elements (*Clock* and *cycle*) in p50T, a bivoltine strain which exhibits photoperiodic diapause induction during both embryonic and larval stages. The temporal expression patterns of clock genes changed in each core clock gene knockout strain, suggesting disruption of normal feedback loops produced by circadian clock genes. Furthermore, photoperiodic diapause induction during both embryonic and larval stages was lost in all knockout strains. Our results indicate the involvement of circadian clock in photoperiodic diapause induction in *B. mori*.

**Highlights:**

- We established knockout *Bombyx mori* strains of four core clock genes
- The temporal expression patterns of clock genes changed in knockout strains
- Photoperiodic diapause induction was not observed in any knockout strains

**Graphical abstract:** 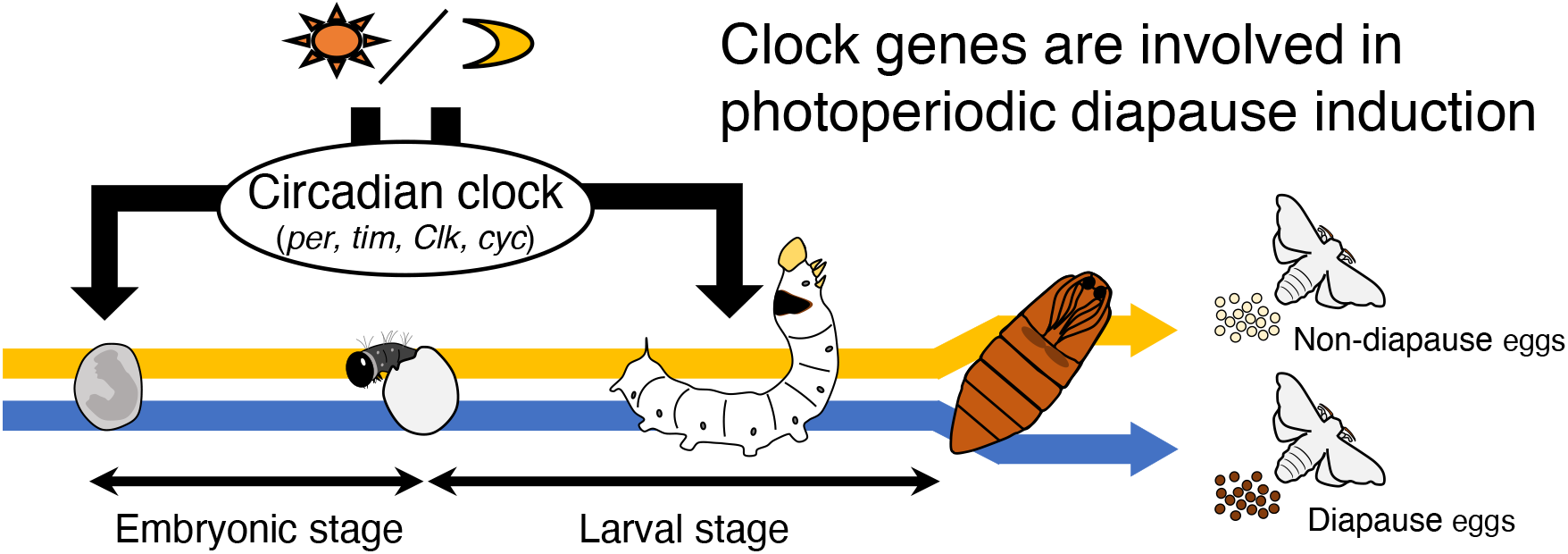

## 1. Introduction

Winter is a severe season for temperate-zone insects because of the cold temperature, drought, and food shortage. To overcome such a difficult season, most insects arrest their development and enter diapause, an inactive physiological state that increases environmental stress-resistance (Denlinger, 1991). Developmental life stages of diapause are diverse among insect species even within orders and sometimes even within genus, suggesting that diapause has evolved multiple times in insects (Meuti and Denlinger, 2013; Nylin, 2013). For example, in Lepidoptera, diapause is induced at the embryonic stage in the silkworm, *Bombyx mori* (Kogure, 1933), at the larval stage in the European corn borer, *Ostrinia nubilalis* (Beck and Hanec, 1960), at the pupal stage in the oak silkworm, *Antheraea pernyi* (Williams and Adkisson, 1964), and at the adult stage in the monarch butterfly, *Danaus plexippus* (Herman, 1981). While diapause stages are genetically fixed, insects can properly time diapause thanks to environmental information, such as the photoperiod and temperature (Denlinger et al., 2012). Although the endocrine control of diapause induction is well known (Denlinger et al., 2012), how insects accept those environment signals, translate the information, and timely activate the endocrine system is not fully understood.

In widely distributed insect species, there is geographical variation in the critical photoperiod (the incidence of diapause is 50% of its maximal level) for diapause induction (Bradshaw and Holzapfel, 1975; Bradshaw and Lounibos, 1977). The critical photoperiod generally increases with latitude among populations of the same species, because of longer summer day length and earlier winter arrival in higher latitudes. Several reports indicate that the critical photoperiod variation is caused by polymorphism in circadian clock genes (Paolucci et al., 2016; Pruisscher et al., 2018; Yamada and Yamamoto, 2011). Clock genes control rhythmic gene expression through autoregulatory feedback loops producing circadian rhythms (Hardin, 2005). These autoregulatory mechanisms are well studied in the fruit fly, *Drosophila melanogaster* (Beer and Helfrich-Förster, 2020). The protein products of the clock genes, *Clock* (*Clk*) and *cycle* (*cyc*), heterodimerize with each other activating the transcription of *period* (*per*) and *timeless* (*tim*) (Allada et al., 1998; Rutila et al., 1998). Then, their protein products, PER and TIM, heterodimerize with each other, translocate to the nucleus, and suppress their own transcriptional activation by inhibiting CLK/CYC (Darlington et al., 1998). This core feedback loop produces the rhythmic expression of *per*, *tim*, and clock-controlled genes that mediate downstream signaling cascades. It is also known that other circadian clock genes, such as *clockwork orange* (*cwo*), *Par-domain protein 1* (*Pdp1*), and *vrille* (*vri*) participate in the core feedback loop producing interlocked feedback loops (Cyran et al., 2003; Glossop et al., 2003; Kadener et al., 2007; Lim et al., 2007; Matsumoto et al., 2007). In addition, TIM is degraded by blue-light dependent binding of the photoreceptor CRYPTOCHROME (CRY) (Ceriani et al., 1999). Therefore, PER/TIM heterodimerization and subsequent CLK/CYC suppression only occur at night. Although the essential features are conserved among insects, the circadian clock components are slightly different (Sandrelli et al., 2008; Tomioka and Matsumoto, 2015). For example, almost all non-drosophilid insects possess *mammalian type cry* (*cry-m* or *cry2*), which has no photoreception ability, in addition to *Drosophila* type *cry* (*cry-d* or *cry1*) (Sandrelli et al., 2008; Tomioka and Matsumoto, 2015). In the monarch butterfly, *Danaus plexippu*, CRY2 functions as a negative element by forming a complex with PER and TIM to inhibit CLK/CYC-mediated transcription (Zhu et al., 2008).

Reduction or loss of core clock genes is reported to perturb normal photoperiodic diapause induction in insects (Table S1). However, it is difficult to distinguish whether the circadian clock as a whole or individual clock genes are responsible for photoperiodic diapause induction (Emerson et al., 2009). To investigate this, the effects of the loss-of-function of multiple single clock genes on photoperiodic diapause induction have been evaluated in the past. In the bean bug, *Riptortus pedestris*, the involvement of core clock genes *cry-m* (*cry-2*), *per, Clk*, and *cyc* in photoperiodic regulation of ovarian development (*i.e*., reproductive diapause) was investigated with RNA interference (RNAi) (Ikeno et al., 2010, 2011, 2013). In the Northern house mosquito, *Culex pipiens*, RNAi against negative elements, *cry2, per*, and *tim* averted reproductive diapause phenotypes under diapause-inducing conditions. In addition, in *D. plexippus*, genetic knockouts by genome editing tools revealed that *Clk, cyc*, and *cry2* are necessary for the photoperiodic responses of oocyte maturation (Iiams et al., 2019). These reports support that circadian clock feedback loops participate in photoperiodic diapause induction in various insect orders. However, since diapause has evolved multiple times in insects, it is noteworthy to examine whether the involvement of circadian clock genes in diapause induction is widely conserved using insects that enter diapause in different stages.

The embryonic diapause of *B. mori* is induced by a maternal diapause hormone, synthesized in the suboesophageal ganglion and secreted from the corpus cardiacum into the hemolymph during the pupal stage(Sato et al., 1998; Takeda and Ogura, 1976). In bivoltine strains, its secretion is determined by environmental conditions, such as temperature and photoperiod. Temperatures >21°C during the late embryonic stage are sensed by a thermoreceptor, transient receptor potential ankyrin 1 (TRPA1), which suppresses diapause hormone secretion (Sato et al., 2014). It is unclear how the “thermal-memory” during the late embryonic stage suppresses diapause hormone secretion during the pupal stage, but the cerebral γ-aminobutyric acid (GABA) ergic and corazonin pathways modulate diapause hormone release via temperature-dependent expression of a plasma membrane GABA transporter (Tsuchiya et al., 2020). Diapause induction in *B. mori* is also affected by the photoperiod during the late embryonic and late larval stages. Diapause induction is promoted by long-day conditions during the late embryonic stage and short-day conditions during the late larval stage (Kogure, 1933). Recently, *per* and *tim* knockout mutants were generated using the genome editing tools TALENs and CRISPR/Cas9, respectively (Ikeda et al., 2019; Nartey et al., 2021). Both mutants lost eclosion and hatching rhythms, indicating that core clock genes are indispensable for circadian rhythms in *B. mori*. In addition, two groups independently described *per’s* involvement in photoperiodic diapause induction (Cui et al., 2021; Ikeda et al., 2021). Ikeda et al. demonstrated that *per* knockout in the Kosetsu strain abolishes photoperiod sensitivity during the larval stage and inhibits diapause egg production. Cui et al. (2021) showed knocking out *per* in the Dazao strain attenuates the effects of temperature and photoperiod on diapause induction during the embryonic stage thorough GABAergic signals. Since both studies only focused on a single negative element, it is difficult to conclude whether clock genes control photoperiodic diapause induction as a unit. In this study, we performed CRISPR/Cas9-mediated gene knockouts of both negative, *per* and *tim*, and positive elements, *Clk* and *cyc*, using a bivoltine strain and investigated the effects on photoperiodic diapause induction at both embryonic and larval stages.

## 2. Materials and methods

### 2.1. Silkworm strains

A bivoltine inbred strain p50T (Daizo) was used in this study. The strain was maintained in our laboratory with mulberry leaves or artificial diet SilkMate PS (NOSAN) under a daily 12 h light:12 h dark (12L12D) cycle at 25°C.

### 2.2. CRISPR/Cas9-mediated gene knockouts

CRISPR/Cas9 was used to knockout *B*. *mori* core clock genes. A unique single guide RNA (sgRNA) was designed for the target genes, *per* (gene model, KWMTBOMO00426), *tim* (gene model, KWMTBOMO01950), *Clk* (full-length cDNA clone, AK380522), and *cyc* (gene model, KWMTBOMO00654) using CRISPRdirect (Naito et al., 2015: https://crispr.dbcls.jp/) based on their coding sequence. sgRNA synthesis was conducted according to the method of Bassett et al. (2014) using the primers listed in Table S2. Cas9 protein (600 ng/μL; NIPPON GENE) and sgRNA (150 ng/μL) were mixed in injection buffer (100 mM KOAc, 2 mM Mg (OAc)2, 30 mM HEPES-KOH; pH 7.4) and injected into non-diapause eggs within 2–4 h after oviposition.

### 2.3. Establishment of knockout strains

Hatched G_0_ (injected generation) larvae were raised to adulthood and crossed with wild-type moths to obtain G_1_ eggs. Genomic DNA was briefly extracted from a leg of each G_1_ adult moth by the HotSHOT method (Truett et al., 2000). DNA fragments containing sgRNA target sequence were amplified by genomic PCR using KOD One (TOYOBO) with the primers listed in Table S2. The PCR reaction was conducted as follows: 35–40 cycles at 98°C for 10 s, 60°C for 5 s, and 68°C for 5 s/kb. Mutations were detected by Heteroduplex Mobility Assay using the MultiNA (SHIMADZU) microchip electrophoresis system (Ansai et al., 2014; Ota et al., 2013). G_1_ moths with an identical mutation were intercrossed with each other to generate a homozygous knockout strain. PCR products were also sequenced using the ABI3130xl genetic analyzer (Applied Biosystems) to confirm the mutation sequence.

### 2.3. Quantitative real-time PCR

Non-diapause eggs were maintained at 25°C under continuous darkness (DD) until hatching. Hatched larvae were reared with artificial diets under 12L12D during the larval stage. Day 3 5th-instar larvae were instantly frozen at Zeitgeber time (ZT) 1, ZT 5, ZT 9, ZT 13, ZT 17, and ZT 21 under 12L12D conditions and stored at −80°C until use. Total RNA was extracted from a whole head using TRIzol reagent (Invitrogen) and then used to synthesize cDNA with TaKaRa RNA PCR™ Kit (AMV) Ver.3.0 (TaKaRa). Quantitative real-time PCR (qPCR) was carried out using a KAPA™ SYBR FAST qPCR Kit (KAPA Biosystems) and StepOnePlus™ Real-Time PCR System (Applied Biosystems), and the transcript levels of core clock genes were calculated by the 2^-ΔΔCt^ method.*actin A3* and *A4* genes were used as reference to normalize the mRNA level of each gene. The primers used for qPCR are listed in Table S2.

One-way ANOVA was used to examine the statistical significance of temporal differences in clock gene expression in each strain. Two-way ANOVA was used to detect the statistical significance of differences in clock gene expression between wild-type and each knockout strain.

### 2.4. Diapause phenotyping

For diapause phenotyping, non-diapause eggs from all strains were used; larvae were fed with artificial diet SilkMate PS (NOSAN) to arrange nutritional conditions. To verify the effects of clock gene knockouts on photoperiodic diapause induction during the larval stage, eggs were incubated at 25°C under DD and the hatched larvae were reared under long-day (non-diapause) (20L4D) or short-day (diapause) conditions (8L16D). On the other hand, to verify the effects during the embryonic stage, eggs were incubated at 18°C under DD or continuous light (LL) until hatching and the larvae reared at 25°C under 12L12D. The resultant females were crossed with males, and the oviposited eggs were maintained at 25°C for 7–8 days. Ommochrome-pigmented (colored) eggs were judged as diapause eggs, whereas uncolored and developed eggs until the head pigmentation stage were considered non-diapause eggs. Uncolored and undeveloped eggs were judged as unfertilized and excluded from the calculation. The ratio of diapause and non-diapause eggs was counted in each batch. When >90% eggs were diapause eggs, the batch was considered as “diapause.” On the other hand, if >90% eggs were non-diapause eggs, the batch was considered as “non-diapause.” When ≥10% eggs were mixed in the same batch, the batch was considered as “mix.” Then, the diapause phenotype of each strain was evaluated by the ratio of diapause, non-diapause, and mix batches to the total batch number.

## 3. Results and discussion

### 3.1 Establishment of core clock gene knockout strains

To investigate the involvement of circadian clock in photoperiodic diapause induction in *B. mori*, we knocked out four core clock genes using CRISPR/Cas9. The inbred strain p50T was used in this study, because our group already published high-quality genome assembly and predicted gene models on this strain (Kawamoto et al., 2019). In addition, the original bivoltine strain “Daizo” exhibits photoperiodic diapause induction during both embryonic and larval stages (Shimizu and Hasegawa, 1988; Egi et al., 2014). We designed sgRNAs targeting the 7th and 6th exon of the negative elements, *per* and *tim*, respectively (Fig. 1A, B). We also designed sgRNAs targeting the 7th exon of the positive elements, *Clk* and *cyc* (Fig. 1C, D). We injected each sgRNA with Cas9 protein into non-diapause eggs and successfully established four core clock gene knockout strains. In each strain, a premature stop codon was generated by site specific random mutations produced by non-homologous end joining and a frame shift, resulting in C-terminally truncated proteins. As all truncated proteins lacked some functional domains, protein function was probably disrupted in all knockout strains.

**Figure 1.**
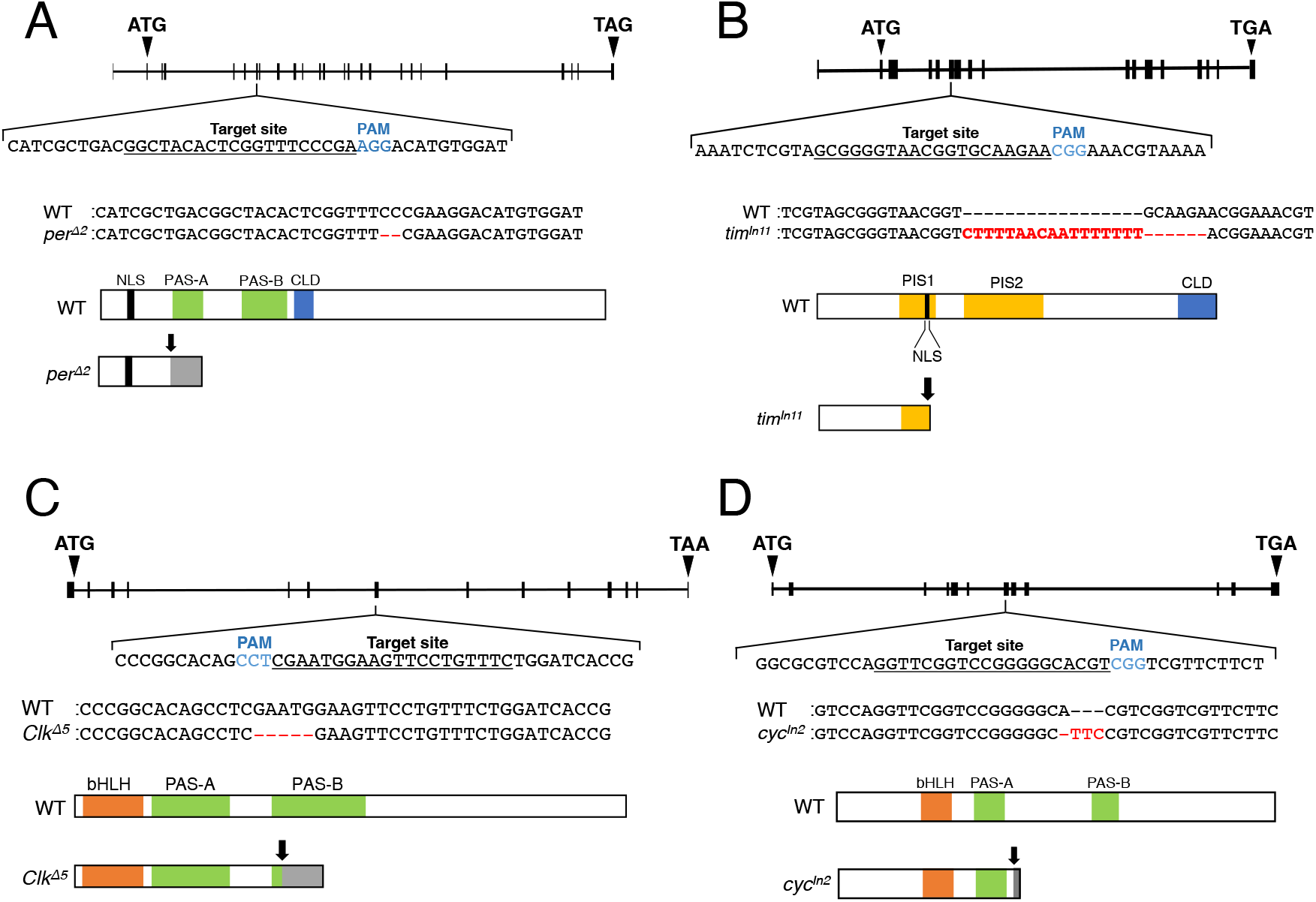
CRISPR/Cas9-mediated knockouts of core clock genes. Genomic structures of *period* (A), *timeless* (B), *Clock* (C), and *cycle* (D) with the sgRNA target site (underlined) and the protospacer adjacent motif (PAM) site (blue letters) at the top. Alignments of nucleotide sequences surrounding the sgRNA target site from wild-type (WT) and knockout strains at the center. Mutations are indicated by red letters. Predicted protein structures are shown at the bottom. Black arrows indicate the mutation site of each clock gene. Functional domains are shown by colored boxes. Nuclear localization signal (NLS; black), PER-ARNT-SIM (PAS; green), cytoplasmic localization domain (CLD; blue), PER interaction site (PIS; yellow), and basic helix-loop-helix (bHLH; orange).

The negative element PER possesses two PER-ARNT-SIM (PAS) domains (PAS-A and PAS-B) involved in heterodimerization with TIMELESS and cytoplasmic localization domain (CLD, Fig. 1A; Iwai et al., 2006). The *per^Δ2^* strain had a 2-bp deletion in the targeting site of exon 7 and potentially produced truncated PER protein lacking the two PAS domains and the CLD. Ikeda et al. (2019, 2021) also established two *per* knockout strains using TALENs targeting the same exon and observed circadian rhythm disruption. On the other hand, Cui et al. (2021) targeted just downstream of the translation start site to establish a *per* knockout strain using TALENs.

Another negative element, TIM, possesses two PER interaction sites (PIS) and CLD (Fig. 1B; Iwai et al., 2006). The *tim^In11^* strain had a 6-bp deletion and a 17-bp insertion in the targeting site of exon 6 and potentially produced a truncated protein lacking a part of PIS1, the whole PIS2, and the C-terminal CLD. Nartey et al. (2021) obtained *tim* knockout mutants by CRISPR/Cas9 targeting two sgRNA target sites located in adjacent exons and observed circadian rhythm disruption.

The positive element CLK possess a basic helix-loop-helix (bHLH) domain and two PAS domains involved in heterodimerization with CYC (Fig. 1C; Allada et al., 1998). The *Clk^Δ5^* strain lacked most of the PAS-B domain due to a premature stop codon generated by a 5-bp deletion in the target site of exon 7.

CYC, another positive element, has a bHLH domain and two PAS domains (Fig. 1D; Markova et al., 2003). The *cyc^In2^* strain lacked the whole PAS-B domain due to a premature stop codon generated by a 1-bp deletion and 3-bp insertion in the target site of exon 7.

### 3.2 Expression analysis of core clock genes

The temporal expression patterns of core clock genes under 12L12D were examined by qPCR using heads of 5th-instar larvae on the 3rd day after ecdysis (Fig. 2, 3). In the wild-type strain, weak but clear temporal changes in the expression of the negative elements, *per* and *tim*, were observed, although there were no significant differences in *tim* expression in two of the four trials (*per*: *P* < 0.001, *tim*: 0.0133 ≤ *P* ≤ 0.242, one-way ANOVA). Ikeda et al. (2021) also analyzed *per* and *tim* expression under almost the same conditions using another bivoltine strain, Kosetsu. However, the expression patterns of both negative elements differed between p50T and Kosetsu. This may be caused by differences in *B. mori* strains since previous studies showed differing *per* and *tim* expression rhythms among tissues, stages, and strains (Cui et al., 2021; Ikeda et al., 2019, 2021; Iwai et al., 2006; Tao et al., 2017). In the *per^Δ2^* strain, *tim* expression was upregulated at all time points (*P* < 0.001, two-way ANOVA; Fig. 2A). This result indicates that knocking out *per* increased *tim’s* transcriptional level through the lack of negative feedback. In other words, this result suggests that PER is involved in the repression of *tim’s* CLK/CYC-mediated transcriptional activation in *B. mori*. On the other hand, although *tim* mRNA levels decreased in *tim^In11^* strain (*P* < 0.001, two-way ANOVA), knocking out *tim* did not increase *per’s* transcriptional levels (*P* = 0.3835, two-way ANOVA; Fig. 2B). This might be due to the loss of CLK/CYC-mediated transcriptional activation in the p50T strain, as described below.

**Figure 2.**
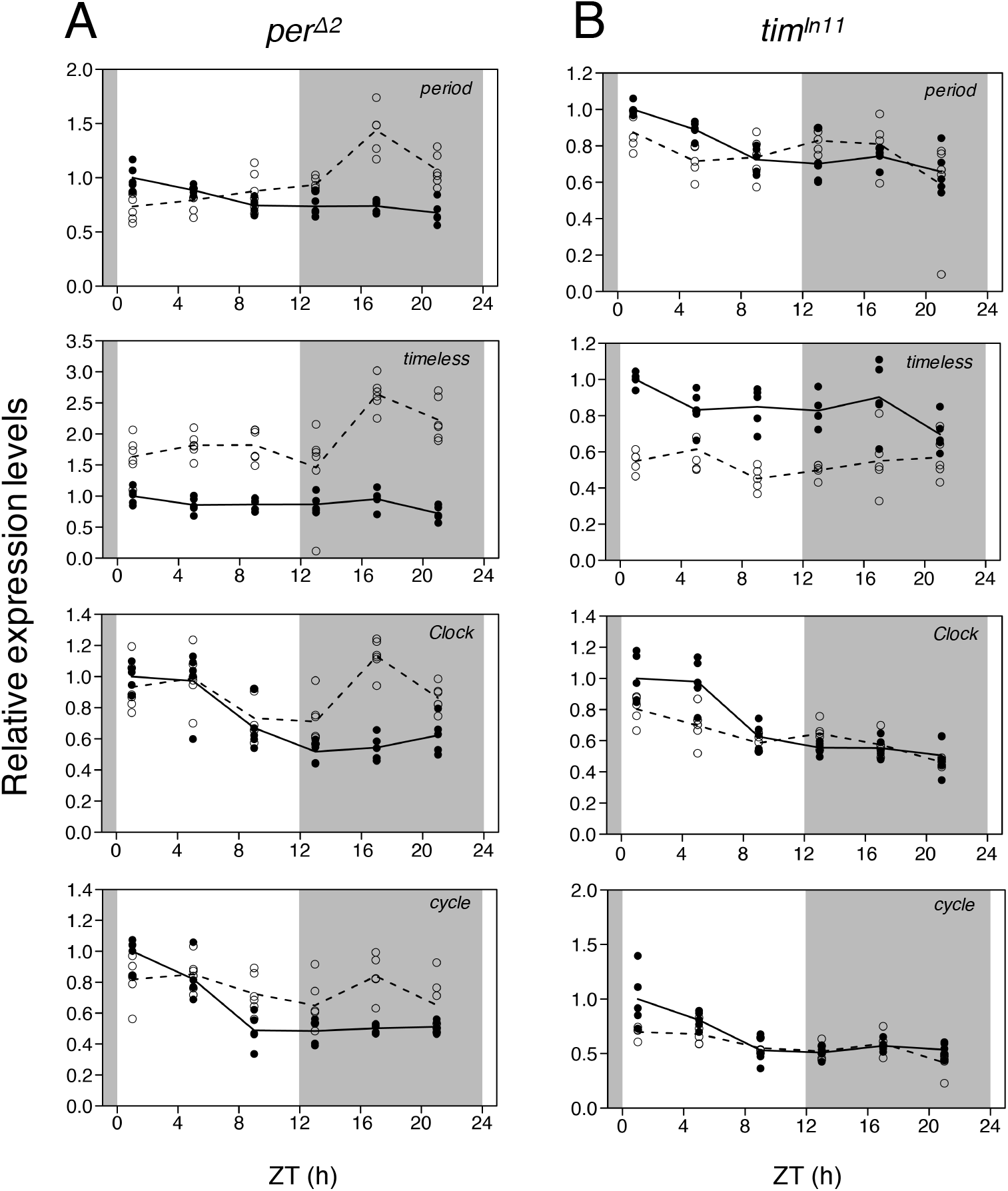
Temporal expression patterns of core clock genes in knockout strains of negative elements. The expression levels of *per*, *tim, Clk*, and *cyc* in heads of day 3 5th-instar larvae under 12L12D were examined by qPCR. The relative expression levels (the means of wild-type at Zeitgeber time (ZT) 1 = 1) of wild-type (closed circles) and knockout strains (open circles), *per^Δ2^* (A) and *tim^In11^* (B), are shown by solid and broken lines, respectively (n = 5 or 6). *actin A3* and *A4* genes were used as reference to normalize the mRNA level of each clock gene. Shaded areas indicate scotophase.

**Figure 3.**
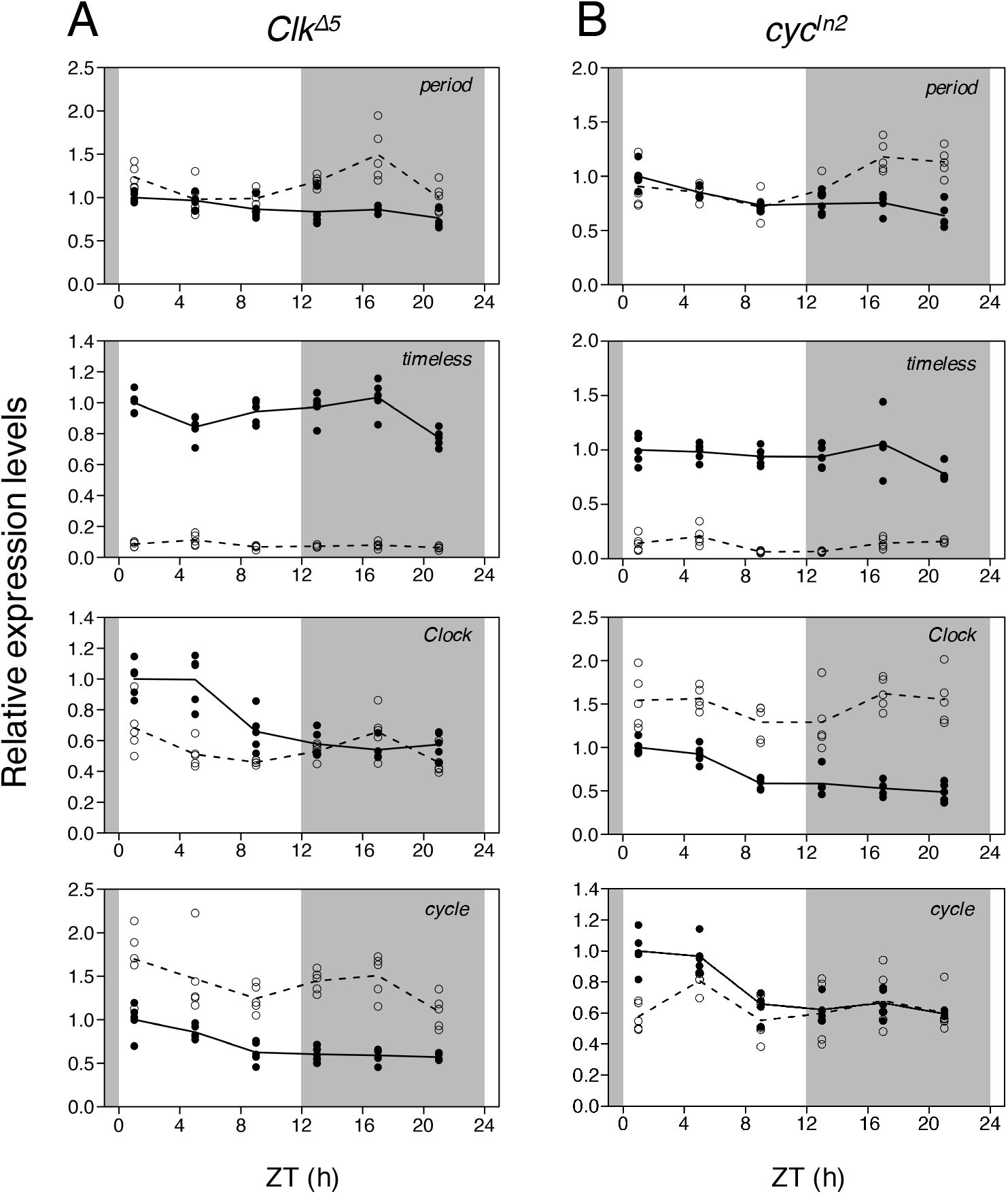
Temporal expression patterns of core clock genes in knockout strains of positive elements. The expression levels of *per*, *tim, Clk*, and *cyc* in heads of day 3 5th-instar larvae under 12L12D were examined by qPCR. The relative expression levels (the means of wild-type at Zeitgeber time (ZT) 1 = 1) of wild-type (closed circles) and knockout strains (open circles), *Clk^Δ5^* (A) and *cyc^In2^* (B), are shown by solid and broken lines, respectively (n = 5). *actin A3* and *A4* genes were used as reference to normalize the mRNA level of each clock gene. Shaded areas indicate scotophase.

*Clk* and *cyc*, positive elements of circadian clock, showed rhythmic expression peaking in the early photophase in the wild-type strain (*P* < 0.001, one-way ANOVA; Fig. 2, 3). The expression patterns of clock genes differ among insect species (Tomioka and Matsumoto, 2015). In *D. melanogaster*, *Clk* is expressed rhythmically, whereas *cyc* expression is constant (Rutila et al., 1998). On the other hands, in honeybees, aphids, crickets, and firebrats, *Clk* is constitutively expressed and *cyc* rhythmically expressed (Cortés et al., 2010; Kamae et al., 2010; Moriyama et al., 2012; Rubin et al., 2006; Uryu et al., 2013). Interestingly, both *Clk* and *cyc* exhibit rhythmic expression patterns in sandflies and mosquitoes (Meireles-Filho and Kyriacou, 2013; Tomioka and Matsumoto, 2015).

In both *Clk^Δ5^* and *cyc^In2^* strains, *tim* mRNA expression levels were extremely low (*P* < 0.001, two-way ANOVA), while *per* expression was not downregulated, but rather upregulated during the scotophase (*P* < 0.001, two-way ANOVA; Fig. 3A, B). These results differ from previous studies in *D. melanogaster* and *D. plexippus*, which showed extremely low mRNA levels of *tim* and *per* mRNA in loss-of-function mutants of positive elements (Allada et al., 1998; Markert et al., 2016; Rutila et al., 1998). Our results suggest that CLK/CYC is involved in transcriptional activation of *tim*, but not *per*. However, it is unclear whether this is a common feature within *B. mori* strains, as this is the first time that *per* and *tim* expression was examined in knockout strains of positive elements.

*Clk* and *cyc* expression levels were significantly higher in *cyc^In2^* and *Clk^Δ5^* strains, respectively (*P* < 0.001, two-way ANOVA; Fig. 3A, B). In *D. melanogaster*, CLK/CYC activates *Pdp1* and *vri* transcription; subsequently, the resultant PDP1 and VRI proteins activate and repress *Clk* transcription, respectively (Cyran et al., 2003; Glossop et al., 2003). The *cyc* and *Clk* upregulation in knockout strains *Clk^Δ5^* and *cyc^In2^* suggests the presence of interlocked feedback loops via *vri* like in *D. melanogaster* and the fall armyworm *Spodoptera frugiperda* (Hänniger et al., 2017; Rego et al., 2021).

### 3.3 Effects of core clock gene knockout on photoperiodic diapause induction

The inbred strain p50T exhibits clear photoperiodic diapause induction during both the embryonic and larval stages (Egi et al., 2014). Using p50T-derived knockout strains, we examined the involvement of core clock genes in photoperiodic diapause induction at each stage. When wild-type larvae were reared under the short-day condition (8L16D) at 25°C, all female moths produced diapause eggs (Fig. 4). On the other hand, under the long-day condition (20L4D), they all produced non-diapause eggs. All female moths produced non-diapause eggs in all core clock gene knockout strains, *per^Δ2^, tim^In11^, Clk^Δ5^*, and *cyc^In2^*, regardless of photoperiodic conditions (Fig. 4). Our results agree with Ikeda et al. (2021) that found that knocking out *per* in Kosetsu, another bivoltine strain, inhibits the diapause egg production induced by a short-day condition during the larval stage. Recently, Homma et al. (2022) reported that core clock genes regulate temperature-dependent diapause induction in *B*. *mori* using TALEN-mediated knockouts in the Kosetsu strain. Although the hatching condition in this experiment (25°C DD) is also affected by temperature-dependent diapause induction during the embryonic stage, our results support that core clock genes are involved in photoperiodic diapause induction during the larval stage.

**Figure 4.**
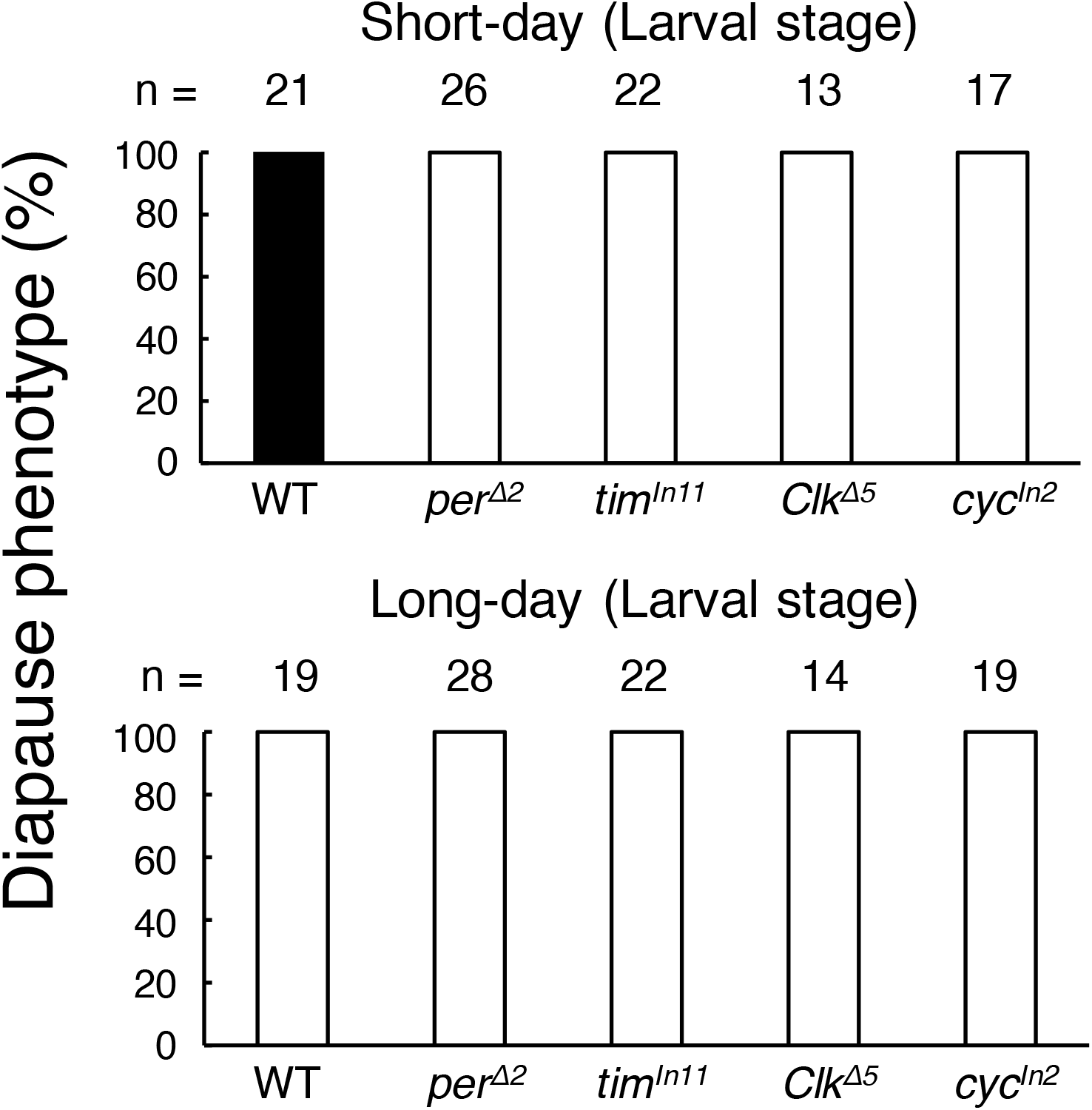
Effects of core clock gene knockouts on photoperiodic diapause induction during larval stages. Larvae were reared under short-day (8L16D; top) or long-day (20L4D; bottom) conditions. Closed and open bars show the proportions of females that oviposited diapause and non-diapause eggs, respectively. No female oviposited both diapause and non-diapause eggs in the same batch.

Next, we examined the involvement of core clock genes in photoperiodic diapause induction during embryonic stages. Under low temperature (18°C), continuous light during the embryonic period induced diapause egg production in all wild-type females, whereas continuous darkness decreased the number of female moths which produced diapause eggs to 40% (Fig. 5). When the eggs of all core clock gene knockout strains, *per^Δ2^, tim^In11^*, *Clk^Δ5^*, and *cyc^In2^* were maintained under continuous light or darkness conditions, knockout females only produced non-diapause eggs. This means that the photoperiodic diapause induction was disrupted by knocking out core clock genes. Cui et al. (2021) also established a *per* knockout in the Dazao strain and found that lack of PER protein suppressed the diapause egg production induced by continuous light condition during the embryonic stage.

**Figure 5.**
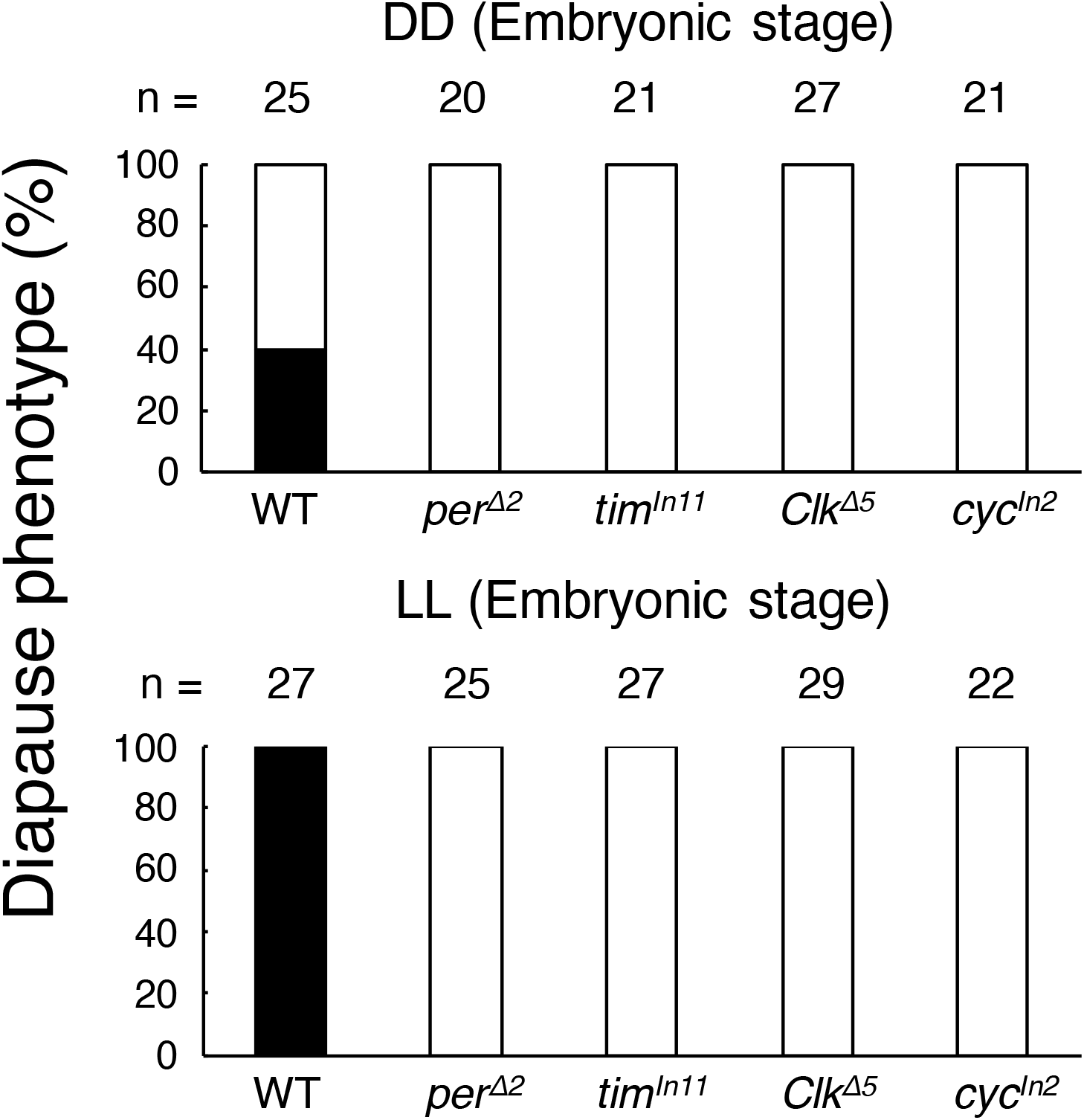
Effects of core clock gene knockouts on photoperiodic diapause induction during embryonic stages. Eggs were incubated under continuous darkness (DD; top) or continuous light (LL; bottom) conditions until hatching. Closed and open bars show the proportions of females that oviposited diapause and non-diapause eggs, respectively. No female oviposited both diapause and non-diapause eggs in the same batch.

Our results together with previous evidence demonstrate that feedback loops involving core clock genes, that is, the circadian clock, participate in photoperiodic diapause induction in *B. mori*. In the monarch butterfly, *D. plexippus*, reproductive diapause is induced by short-day conditions during the larval stage (Goehring and Oberhauser, 2002); the photoperiodic diapause induction is abolished in *Clk* or *cyc* loss-of-function mutant females. In contrast, the reproductive diapause is strongly induced in loss-of-function mutant females for *cry2*, regardless of day length. Although diapause stage and the endocrine system controlling diapause differ between both lepidopteran species, core clock genes are necessary for photoperiodic diapause induction in both. In the bean bug *R. pedestris*, RNAi of the negative elements, *cry-m* (*cry2*) and *per*, causes the insect to avert reproductive diapause under a diapause-inducing photoperiod (short-day) whereas RNAi of the positive elements, *Clk* and *cyc*, induces reproductive diapause under a diapause-averting photoperiod (long-day). These results indicate that circadian activators and repressors may function in a coordinated fashion to sense the photoperiod, probably through a feedback loop. In contrast to previous results, *B. mori* females bearing non-functional positive and negative elements constantly produced non-diapause eggs. These differences among insect species after clock gene perturbation strongly support a conserved involvement of circadian clock feedback loop in photoperiodic diapause induction, despite its independent evolution in each insect species. Since circadian clocks control diverse physiological process, such as behavior, learning, feeding, metabolism, chemosensation, and immunity (Allada and Chung, 2009), we cannot conclude whether circadian clock or non-clock function of clock genes is related to photoperiodic diapause induction in *B. mori*. However, temperature-dependent diapause induction can be rescued by injection of picrotoxin, an ionotropic GABA receptor antagonist, and diapause hormone into female pupae, suggesting that the GABAergic and diapause hormone signaling pathways are intact in clock gene knockouts (Homma et al., 2022). Since diapause hormone release from the corpus cardiacum is suppressed by GABAergic signaling pathway (Homma et al., 2022; Shimizu et al., 1989), our next goal is to reveal how circadian clock genes are influenced by the photoperiod and how the circadian clock controls the GABAergic signaling pathway and diapause hormone secretion.

## Acknowledgments

We thank the Institute for Sustainable Agro-ecosystem Services, The University of Tokyo, for facilitating the mulberry cultivation and the Biotron Facility at the University of Tokyo for rearing the silkworms. We also thank Dr. Susumu Katsuma for useful advice. We are grateful to Wakako Saito and Natsuki Nakashima for technical assistance for silkworm maintenance. This study was supported by JSPS KAKENHI grant numbers JP17H05047, JP20H02997 to TK and JST SPRING (JPMJSP2108) to HT.

## Author contributions

TK supervised and designed this study. HT conducted most of the experiments. TK and HT analyzed the data and wrote the manuscript.

## Supplementary tables

**Table S1.**
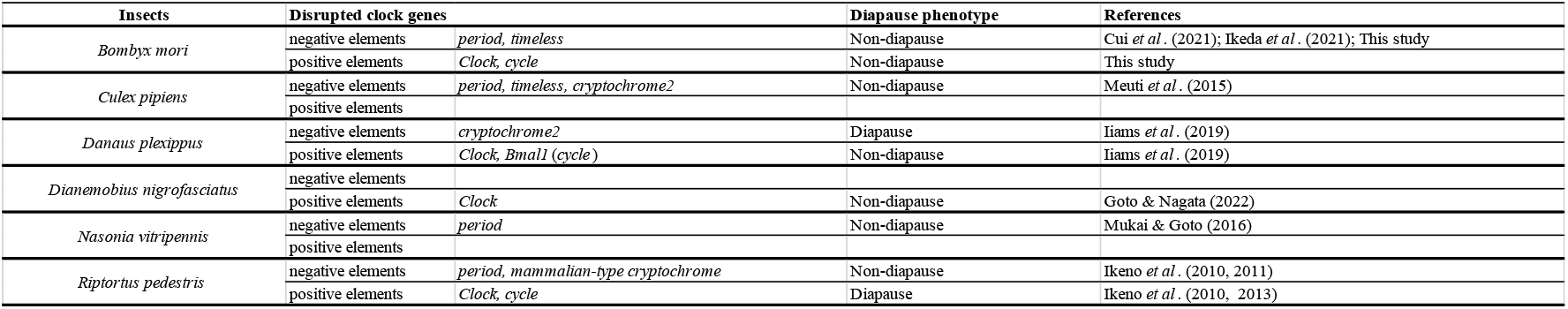
Summary of results of clock gene disruption in insects and its effects on photoperiodic diapause induction.

**Table S2.**
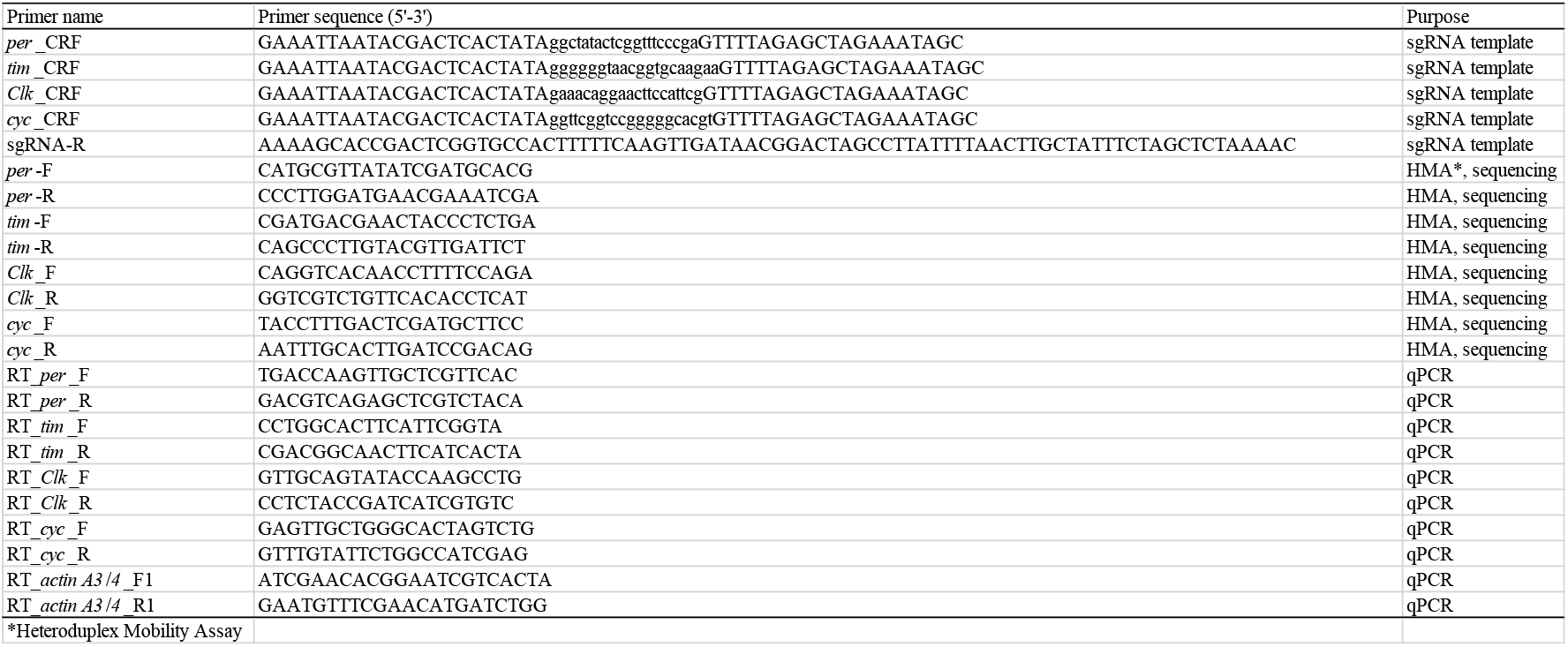
List of primers.

## References

Allada, R., Chung, B.Y., 2009. Circadian organization of behavior and physiology in *Drosophila*. Annu. Rev. Physiol. 72, 605–624. https://doi.org/10.1146/annurev-physiol-021909-135815

Allada, R., White, N.E., So, W.V., Hall, J.C., Rosbash, M., 1998. A mutant *Drosophila* homolog of mammalian *Clock* disrupts circadian rhythms and transcription of *period* and *timeless*. Cell 93, 791–804. https://doi.org/10.1016/S0092-8674(00)81440-3

Ansai, S., Inohaya, K., Yoshiura, Y., Schartl, M., Uemura, N., Takahashi, R., Kinoshita, M., 2014. Design, evaluation, and screening methods for efficient targeted mutagenesis with transcription activator-like effector nucleases in medaka. Dev. Growth Differ. 56, 98–107. https://doi.org/10.1111/dgd.12104

Bassett, A.R., Tibbit, C., Ponting, C.P., Liu, J.L., 2014. Highly Efficient Targeted Mutagenesis of *Drosophila* with the CRISPR/Cas9 System. Cell Rep. 6, 1178–1179. https://doi.org/10.1016/j.celrep.2014.03.017

Beck, S.D., Hanec, W., 1960. Diapause in the European corn borer, *Pyrausta nubilalis* (Hübn.). J. Insect Physiol. 4, 304–318. https://doi.org/10.1016/0022-1910(60)90056-1

Beer, K., Helfrich-Förster, C., 2020. Model and Non-model Insects in Chronobiology. Front. Behav. Neurosci. 14, 1–23. https://doi.org/10.3389/fnbeh.2020.601676

Bradshaw, W.E., Holzapfel, C.M., 1975. Biology of tree-hole mosquitoes: photoperiodic control of development in northern *Toxorhynchites rutilus* (Coq.). Can. J. Zool. 53, 889–893.

Bradshaw, W.E., Lounibos, L.P., 1977. Evolution of dormancy and its photoperiodic control in pitcher-plant mosquitoes. Evolution (N. Y). 31, 546–567. https://doi.org/10.1111/j.1558-5646.1977.tb01044.x

Ceriani, M.F., Darlington, T.K., Staknis, D., Más, P., Petti, A.A., Weitz, C.J., Kay, S.A., 1999. Light-dependent sequestration of TIMELESS by CRYPTOCHROME. Science. 285, 553–556. https://doi.org/10.1126/science.285.5427.553

Cortés, T., Ortiz-Rivas, B., Martínez-Torres, D., 2010. Identification and characterization of circadian clock genes in the pea aphid *Acyrthosiphon pisum*. Insect Mol. Biol. 19, 123–139. https://doi.org/10.1111/j.1365-2583.2009.00931.x

Cui, W.Z., Qiu, J.F., Dai, T.M., Chen, Z., Li, J.L., Liu, K., Wang, Y.J., Sima, Y.H., Xu, S.Q., 2021. Circadian Clock Gene *Period* Contributes to Diapause via GABAeric-Diapause Hormone Pathway in *Bombyx mori*. Biology, 10, 842. https://doi.org/10.3390/biology10090842

Cyran, S.A., Buchsbaum, A.M., Reddy, K.L., Lin, M.-C., Glossop, N.R.J., Hardin, P.E., Young, M.W., Storti, R. V, Blau, J., 2003. *vrille, Pdp1*, and *dClock* Form a Second Feedback Loop in the *Drosophila* Circadian Clock. Cell 112, 329–341.

Darlington, T.K., Wager-Smith, K., Ceriani, M.F., Staknis, D., Gekakis, N., Steeves, T.D.L., Weitz, C.J., Takahashi, J.S., Kay, S.A., 1998. Closing the circadian loop: CLOCK-induced transcription of its own inhibitors *per* and *tim*. Science. 280, 1599–1603. https://doi.org/10.1126/science.280.5369.1599

Denlinger, D.L., 1991. Relationship between Cold Hardiness and Diapause, in: Lee, R.E., Denlinger, D.L. (Eds.), Insects at Low Temperature. Springer, Boston, MA, pp. 174–198.

Denlinger, D.L., Yocum, G.D., Rinehart, J.P., 2012. Hormonal Control of Diapause, in: Gilbert, L.I. (Ed.), Insect Endocrinology. Academic Press, San Diego, pp. 430–463.

Egi, Y., Akitomo, S., Fujii, T., Banno, Y., Sakamoto, K., 2014. Silkworm strains that can be clearly destined towards either embryonic diapause or direct development by adjusting a single ambient parameter during the preceding generation. Entomol. Sci. 17, 396–399. https://doi.org/10.1111/ens.12073

Emerson, K.J., Bradshaw, W.E., Holzapfel, C.M., 2009. Complications of complexity: integrating environmental, genetic and hormonal control of insect diapause. Trends Genet. 25, 217–225. https://doi.org/10.1016/j.tig.2009.03.009

Glossop, N.R.J., Houl, J.H., Zheng, H., Ng, F.S., Dudek, S.M., Hardin, P.E., 2003. VRILLE feeds back to control circadian transcription of *Clock* in the *Drosophila* circadian oscillator. Neuron 37, 249–261. https://doi.org/10.1016/S0896-6273(03)00002-3

Goehring, L., Oberhauser, K.S., 2002. Effects of photoperiod, temperature, and host plant age on induction of reproductive diapause and development time in *Danaus plexippus*. Ecol. Entomol. 27, 674–685. https://doi.org/10.1046/j.1365-2311.2002.00454.x

Hänniger, S., Dumas, P., Schöfl, G., Gebauer-Jung, S., Vogel, H., Unbehend, M., Heckel, D.G., Groot, A.T., 2017. Genetic basis of allochronic differentiation in the fall armyworm. BMC Evol. Biol. 17, 1–14. https://doi.org/10.1186/s12862-017-0911-5

Hardin, P.E., 2005. The circadian timekeeping system of *Drosophila*. Curr. Biol. 15, 714–722. https://doi.org/10.1016/j.cub.2005.08.019

Herman, W.S., 1981. Studies on the Adult Reproductive Diapause of the Monarch Butterfly, *Danaus Plexippus*. Biol. Bull. 160, 89–106. https://doi.org/10.2307/1540903

Homma, S., Murata, A., Ikegami, M., Kobayashi, M., Yamazaki, M., Ikeda, K., Daimon, T., Numata, H., Mizoguchi, A., Shiomi, K., 2022. Circadian Clock Genes Regulate Temperature-Dependent Diapause Induction in Silkworm *Bombyx mori*. Front. Physiol. 13, 1–9. https://doi.org/10.3389/fphys.2022.863380

Iiams, S.E., Lugena, A.B., Zhang, Y., Hayden, A.N., Merlin, C., 2019. Photoperiodic and clock regulation of the vitamin A pathway in the brain mediates seasonal responsiveness in the monarch butterfly. Proc. Natl. Acad. Sci. U. S. A. 116, 25214–25221. https://doi.org/10.1073/pnas.1913915116

Ikeda, K., Daimon, T., Sezutsu, H., Udaka, H., Numata, H., 2019. Involvement of the Clock Gene *period* in the Circadian Rhythm of the Silkmoth *Bombyx mori*. J. Biol. Rhythms 34, 283–292. https://doi.org/10.1177/0748730419841185

Ikeda, K., Daimon, T., Shiomi, K., Udaka, H., Numata, H., 2021. Involvement of the Clock Gene *period* in the Photoperiodism of the Silkmoth *Bombyx mori*. Zoolog. Sci. 38, 523–530. https://doi.org/10.2108/zs210081

Ikeno, T., Ishikawa, K., Numata, H., Goto, S.G., 2013. Circadian clock gene *Clock* is involved in the photoperiodic response of the bean bug *Riptortus pedestris*. Physiol. Entomol. 38, 157–162. https://doi.org/10.1111/phen.12013

Ikeno, T., Numata, H., Goto, S.G., 2011. Photoperiodic response requires *mammalian-type cryptochrome* in the bean bug *Riptortus pedestris*. Biochem. Biophys. Res. Commun. 410, 394–397. https://doi.org/10.1016/j.bbrc.2011.05.142

Ikeno, T., Tanaka, S.I., Numata, H., Goto, S.G., 2010. Photoperiodic diapause under the control of circadian clock genes in an insect. BMC Biol. 8. https://doi.org/10.1186/1741-7007-8-116

Iwai, S., Fukui, Y., Fujiwara, Y., Takeda, M., 2006. Structure and expressions of two circadian clock genes, *period* and *timeless* in the commercial silkmoth, *Bombyx mori*. J. Insect Physiol. 52, 625–637. https://doi.org/10.1016/j.jinsphys.2006.03.001

Kadener, S., Stoleru, D., McDonald, M., Nawathean, P., Rosbash, M., 2007. *Clockwork Orange* is a transcriptional repressor and a new *Drosophila* circadian pacemaker component. Genes Dev. 21, 1675–1686. https://doi.org/10.1101/gad.1552607

Kamae, Y., Tanaka, F., Tomioka, K., 2010. Molecular cloning and functional analysis of the clock genes, *Clock* and *cycle*, in the firebrat *Thermobia domestica*. J. Insect Physiol. 56, 1291–1299. https://doi.org/10.1016/j.jinsphys.2010.04.012

Kawamoto, M., Jouraku, A., Toyoda, A., Yokoi, K., Minakuchi, Y., Katsuma, S., Fujiyama, A., Kiuchi, T., Yamamoto, K., Shimada, T., 2019. High-quality genome assembly of the silkworm, *Bombyx mori*. Insect Biochem. Mol. Biol. 107, 53–62. https://doi.org/10.1016/j.ibmb.2019.02.002

Kogure, M., 1933. The Influence Of Light And Temperature On Certain Characters Of The Silkworm, *Bombyx Mori*. J. Dep. Agric. Kyushu Imp. Univ. 4, 1–93. https://doi.org/10.5109/22568

Lim, C., Chung, B.Y., Pitman, J.L., McGill, J.J., Pradhan, S., Lee, J., Keegan, K.P., Choe, J., Allada, R., 2007. *clockwork orange* Encodes a Transcriptional Repressor Important for Circadian-Clock Amplitude in *Drosophila*. Curr. Biol. 17, 1082–1089. https://doi.org/10.1016/j.cub.2007.05.039

Markert, M.J., Zhang, Y., Enuameh, M.S., Reppert, S.M., Wolfe, S.A., Merlin, C., 2016. Genomic access to monarch migration using TALEN and CRISPR/Cas9-mediated targeted mutagenesis. G3 Genes, Genomes, Genet. 6, 905–915. https://doi.org/10.1534/g3.116.027029

Markova, E.P., Ueda, H., Sakamoto, K., Oishi, K., Shimada, T., Takeda, M., 2003. Cloning of *Cyc* (*Bmal1*) homolog in *Bombyx mori:* Structural analysis and tissue specific distributions. Comp. Biochem. Physiol. - B Biochem. Mol. Biol. 134, 535–542. https://doi.org/10.1016/S1096-4959(03)00004-6

Matsumoto, A., Ukai-Tadenuma, M., Yamada, R.G., Houl, J., Uno, K.D., Kasukawa, T., Dauwalder, B., Itoh, T.Q., Takahashi, K., Ueda, R., Hardin, P.E., Tanimura, T., Ueda, H.R., 2007. A functional genomics strategy reveals *clockwork orange* as a transcriptional regulator in the *Drosophila* circadian clock. Genes Dev. 21, 1687–1700. https://doi.org/10.1101/gad.1552207

Meireles-Filho, A.C.A., Kyriacou, C.P., 2013. Circadian rhythms in insect disease vectors. Mem. Inst. Oswaldo Cruz 108, 48–58. https://doi.org/10.1590/0074-0276130438

Meuti, M.E., Denlinger, D.L., 2013. Evolutionary links between circadian clocks and photoperiodic diapause in insects. Integr. Comp. Biol. 53, 131–143. https://doi.org/10.1093/icb/ict023

Moriyama, Y., Kamae, Y., Uryu, O., Tomioka, K., 2012. *Gb’clock* is expressed in the optic lobe and is required for the circadian clock in the cricket *Gryllus bimaculatus*. J. Biol. Rhythms 27, 467–477. https://doi.org/10.1177/0748730412462207

Naito, Y., Hino, K., Bono, H., Ui-Tei, K., 2015. CRISPRdirect: software for designing CRISPR/Cas guide RNA with reduced off-target sites. Bioinformatics 31, 1120–1123. https://doi.org/10.1093/bioinformatics/btu743

Nartey, M.A., Sun, X., Qin, S., Hou, C.X., Li, M.W., 2021. CRISPR/Cas9-based knockout reveals that the clock gene *timeless* is indispensable for regulating circadian behavioral rhythms in *Bombyx mori*. Insect Sci. 28, 1414–1425. https://doi.org/10.1111/1744-7917.12864

Nylin, S., 2013. Induction of diapause and seasonal morphs in butterflies and other insects: Knowns, unknowns and the challenge of integration. Physiol. Entomol. 38, 96–104. https://doi.org/10.1111/phen.12014

Ota, S., Hisano, Y., Muraki, M., Hoshijima, K., Dahlem, T.J., Grunwald, D.J., Okada, Y., Kawahara, A., 2013. Efficient identification of TALEN-mediated genome modifications using heteroduplex mobility assays. Genes to Cells 18, 450–458. https://doi.org/10.1111/gtc.12050

Paolucci, S., Salis, L., Vermeulen, C.J., Beukeboom, L.W., van de Zande, L., 2016. QTL analysis of the photoperiodic response and clinal distribution of *period* alleles in *Nasonia vitripennis*. Mol. Ecol. 25, 4805–4817. https://doi.org/10.1111/mec.13802

Pruisscher, P., Nylin, S., Gotthard, K., Wheat, C.W., 2018. Genetic variation underlying local adaptation of diapause induction along a cline in a butterfly. Mol. Ecol. 27, 3613–3626. https://doi.org/10.1111/mec.14829

Rego, N. de F.C., Chahad-Ehlers, S., Campanini, E.B., Torres, F.R., de Brito, R.A., 2021. VRILLE shows high divergence among Higher Diptera flies but may retain role as transcriptional repressor of *Clock*. J. Insect Physiol. 133. https://doi.org/10.1016/j.jinsphys.2021.104284

Rubin, E.B., Shemesh, Y., Cohen, M., Elgavish, S., Robertson, H.M., Bloch, G., 2006. Molecular and phylogenetic analyses reveal mammalian-like clockwork in the honey bee (*Apis mellifera*) and shed new light on the molecular evolution of the circadian clock. Genome Res. 16, 1352–1365. https://doi.org/10.1101/gr.5094806

Rutila, J.E., Suri, V., Le, M., So, W.V., Rosbash, M., Hall, J.C., 1998. CYCLE is a second bHLH-PAS clock protein essential for circadian rhythmicity and transcription of *Drosophila period* and *timeless*. Cell 93, 805–814. https://doi.org/10.1016/S0092-8674(00)81441-5

Sandrelli, F., Costa, R., Kyriacou, C.P., Rosato, E., 2008. Comparative analysis of circadian clock genes in insects. Insect Mol. Biol. 17, 447–463. https://doi.org/10.1111/j.1365-2583.2008.00832.x

Sato, A., Sokabe, T., Kashio, M., Yasukochi, Y., Tominaga, M., Shiomi, K., 2014. Embryonic thermosensitive TRPA1 determines transgenerational diapause phenotype of the silkworm, *Bombyx mori*. Proc. Natl. Acad. Sci. U. S. A. 111. https://doi.org/10.1073/pnas.1322134111

Sato, Y., Shiomi, K., Saito, H., Imai, K., Yamashita, O., 1998. Phe-X-Pro-Arg-Leu-NH2 peptide producing cells in the central nervous system of the silkworm, *Bombyx mori*. J. Insect Physiol. 44, 333–342. https://doi.org/10.1016/S0022-1910(97)00140-6

Shimizu, I., Hasegawa, K., 1988. Photoperiodic induction of diapause in the silkworm, *Bombyx mori:* location of the photoreceptor using a chemiluminescent paint. Physiol. Entomol. 13, 81–88. https://doi.org/10.1111/j.1365-3032.1988.tb00911.x

Shimizu, I., Matsui, T., Hasegawa, K., 1989. Possible Involvement of GABAergic Neurons in Regulation of Diapause Hormone Secretion in the Silkworm, *Bombyx mori*. Zoolog. Sci. 6, 809–812.

Takeda, S., Ogura, N., 1976. Induction of egg diapause by implantation of corpora cardiaca and corpora allata in *Bombyx mori*. J. Insect Physiol. 22, 941–944. https://doi.org/10.1016/0022-1910(76)90075-5

Tao, H., Li, X., Qiu, J.F., Liu, H.J., Zhang, D.Y., Chu, F., Sima, Y., Xu, S.Q., 2017. The light cycle controls the hatching rhythm in *Bombyx mori* via negative feedback loop of the circadian oscillator. Arch. Insect Biochem. Physiol. 96, 1–14. https://doi.org/10.1002/arch.21408

Tomioka, K., Matsumoto, A., 2015. Circadian molecular clockworks in non-model insects. Curr. Opin. Insect Sci. 7, 58–64. https://doi.org/10.1016/j.cois.2014.12.006

Truett, G.E., Heeger, P., Mynatt, R.L., Truett, A.A., Walker, J.A., Warman, M.L., 2000. Preparation of PCR-quality mouse genomic dna with hot sodium hydroxide and tris (HotSHOT). Biotechniques 29, 52–54. https://doi.org/10.2144/00291bm09

Tsuchiya, R., Kaneshima, A., Kobayashi, M., Yamazaki, M., Takasu, Y., Sezutsu, H., Tanaka, Y., Mizoguchi, A., Shiomi, K., 2020. Maternal GABAergic and GnRH/corazonin pathway modulates egg diapause phenotype of the silkworm *Bombyx mori*. Proc. Natl. Acad. Sci. U. S. A. 118. https://doi.org/10.1073/pnas.2020028118

Uryu, O., Karpova, S.G., Tomioka, K., 2013. The clock gene *cycle* plays an important role in the circadian clock of the cricket *Gryllus bimaculatus*. J. Insect Physiol. 59, 697–704. https://doi.org/10.1016/j.jinsphys.2013.04.011

Williams, C.M., Adkisson, P.L., 1964. Physiology of Insect Diapause. XIV. An Endocrine Mechanism for the Photoperiodic Control of Pupal Diapause in the Oak Silkworm, *Antheraea pernyi*. Biol. Bull. 127, 511–525.

Yamada, H., Yamamoto, M., 2011. Association between circadian clock genes and diapause incidence in *Drosophila triauraria*. PLoS One 6. https://doi.org/10.1371/journal.pone.0027493

Zhu, H., Sauman, I., Yuan, Q., Casselman, A., Emery-Le, M., Emery, P., Reppert, S.M., 2008. Cryptochromes define a novel circadian clock mechanism in monarch butterflies that may underlie sun compass navigation. PLoS Biol. 6, 0138–0155. https://doi.org/10.1371/journal.pbio.0060004

